# Leaf-GP: An Open and Automated Software Application for Measuring Growth Phenotypes for Arabidopsis and Wheat

**DOI:** 10.1101/180083

**Authors:** Ji Zhou, Christopher Applegate, Albor Dobon Alonso, Daniel Reynolds, Simon Orford, Michal Mackiewicz, Simon Griffiths, Steven Penfield, Nick Pullen

## Abstract

**Background:** Plants demonstrate dynamic growth phenotypes that are determined by genetic and environmental factors. Phenotypic analysis of growth features over time is a key approach to understand how plants interact with environmental change as well as respond to different treatments. Although the importance of measuring dynamic growth traits is widely recognised, available open software tools are limited in terms of batch processing of image datasets, multiple trait analysis, software usability and cross-referencing results between experiments, making automated phenotypic analysis problematic.

**Results:** Here, we present Leaf-GP (Growth Phenotypes), an easy-to-use and open software application that can be executed on different platforms. To facilitate diverse scientific user communities, we provide three versions of the software, including a graphic user interface (GUI) for personal computer (PC) users, a command-line interface for high-performance computer (HPC) users, and an interactive *Jupyter Notebook* (also known as the iPython Notebook) for computational biologists and computer scientists. The software is capable of extracting multiple growth traits automatically from large image datasets. We have utilised it in *Arabidopsis thaliana* and wheat (*Triticum aestivum*) growth studies at the Norwich Research Park (NRP, UK). By quantifying growth phenotypes over time, we are able to identify diverse plant growth patterns based on a variety of key growth-related phenotypes under varied experimental conditions.

As Leaf-GP has been evaluated with noisy image series acquired by different imaging devices and still produced reliable biologically relevant outputs, we believe that our automated analysis workflow and customised computer vision based feature extraction algorithms can facilitate a broader plant research community for their growth and development studies. Furthermore, because we implemented Leaf-GP based on open Python-based computer vision, image analysis and machine learning libraries, our software can not only contribute to biological research, but also exhibit how to utilise existing open numeric and scientific libraries (including Scikit-image, OpenCV, SciPy and Scikit-learn) to build sound plant phenomics analytic solutions, efficiently and effectively.

**Conclusions:** Leaf-GP is a comprehensive software application that provides three approaches to quantify multiple growth phenotypes from large image series. We demonstrate its usefulness and high accuracy based on two biological applications: (1) the quantification of growth traits for *Arabidopsis* genotypes under two temperature conditions; and (2) measuring wheat growth in the glasshouse over time. The software is easy-to-use and cross-platform, which can be executed on Mac OS, Windows and high-performance computing clusters (HPC), with open Python-based scientific libraries preinstalled. We share our modulated source code and executables (.exe for Windows; .app for Mac) together with this paper to serve the plant research community. The software, source code and experimental results are freely available at https://github.com/Crop-Phenomics-Group/Leaf-GP/releases.

## Background

Plants demonstrate dynamic growth phenotypes that are determined by genetic and environmental factors [1–3]. Phenotypic features such as relative growth rates (RGR), vegetative greenness and other morphological characters are popularly utilised by researchers in order to quantify how plants interact with environmental changes (i.e. GxE) [4–6]. In particular, to assess the growth and development in response to various experimental treatments, growth phenotypes (e.g. leaf area, canopy size and leaf numbers) are considered as key measurements [7–12], indicating the importance of dynamically scoring differences of growth related traits between experiments. To accomplish the above tasks, high quality image-based growth data need to be collected from many biological replicates over time [13,14], which is then followed by manual, semi-automated, or automated trait analysis [15,16]. However, the current bottleneck lies in how to extract meaningful results from our increasing phenotypic data, effectively and efficiently [14,17].

To facilitate the quantification of dynamic growth traits, a range of imaging hardware and software have been developed. We summarise some representative tools as follows:

- LeafAnalyser [18] uses image-processing techniques to measure leaf shape variation as well as record the position of each leaf automatically.
- GROWSCREEN [12] quantifies dynamic seedling growth under altered light conditions.
- GROWSCREEN FLUORO [19] measures leaf growth and chlorophyll fluorescence to detect stress tolerance.
- LemnaGrid [20] integrates image analysis and rosette area modelling to assess genotype effects for *Arabidopsis*.
- Leaf Image Analysis Interface (LIMANI) [21] segments and computes venation patterns of *Arabidopsis* leaves.
- Rosette Tracker [22] provides an open Java-based image analysis solution to evaluate plant-shoot phenotypes to facilitate the understanding of *Arabidopsis* genotype effects.
- PhenoPhyte [23] semi-automates the quantification of various 2D leaf traits through a web-based software application.
- OSCILLATOR [24] analyses rhythmic leaf growth movement using infrared photography combined with wavelet transformation in mature plants.
- HPGA (a high-throughput phenotyping platform for plant growth modelling and functional analysis) [5], which produces plant area estimation and growth modelling and analysis to high-throughput plant growth analysis.
- LeafJ [25] provides an ImageJ plugin to semi-automate leaf shape measurement.
- Integrated Analysis Platform (IAP) [16] is an open framework that performs high-throughput plant phenotyping based on the LemnaTec system.
- Easy Leaf Area [26] uses colour-based feature to differentiate and measure leaves from their background using a red calibration area to replace scale measurement.
- Phytotyping^4D^ [27] employs a light-field camera to simultaneously provide a focus and a depth image so that distance information from leaf surface can be quantified.
- Leaf Angle Distribution Toolbox [28] is a Matlab-based software package for quantifying leaf surface properties via 3D reconstruction from stereo images.
- MorphoLeaf [29] is a plug-in for the Free-D software to perform analysis of morphological features of leaves with different architectures.
- rosettR [30] is a high-throughput phenotyping protocol for measuring total rosette area of seedlings grown in plates.
- A real-time machine learning based classification phenotyping framework [31] can extract leaf canopy to rate soybean stress severity.

While many hardware and software solutions have been created, the threshold for employing the existing tools for measuring growth phenotypes is still relatively high. This is due to many analytic software solutions that are either customised for specific hardware platforms (e.g. LemnaTec), or relied on proprietary or specialised software platforms (e.g. Matlab), restricting the accessibility for smaller or not well-funded laboratories to utilise the existing solutions [22]. Hence, data annotation, phenotypic analysis, and results cross-referencing are still frequently done manually in many laboratories, which is time consuming and prone to errors [21].

Available open software tools are also limited in terms of batch processing, multiple trait analysis, and software usability, making automatic phenotypic analysis problematic [30]. In order to provide an open analytics software solution to serve a broader plant research community, we developed Leaf-GP (Growth Phenotypes), an open-source and easy-to-use software solution that can be easily setup for automated analysis using the community driven Python-based scientific and numeric libraries. After continuous development and testing, Leaf-GP can now extract and compare key growth phenotypes reliably from large image series. Some of the growth-related traits are projected leaf area (mm^2^), leaf perimeter (mm), canopy length and width (mm), leaf canopy area (mm^2^), stockiness (%), compactness (%), leaf numbers and greenness (0-255). We demonstrate its high accuracy and usefulness through experiments using *Arabidopsis thaliana* and *Paragon* wheat (a UK spring wheat variety). The software can be executed on most of the mainstream operating systems with Python and Anaconda distribution preinstalled. More importantly, we followed the open software design strategy, which means our work is expandable and new functions or procedures for other plant species can be easily added.

## Methods

### Applying Leaf-GP to plant growth studies

Figure 1 illustrates how Leaf-GP was applied to quantify growth phenotypes for *Arabidopsis* rosettes and *Paragon* wheat over time. To improve the software flexibility, Leaf-GP was designed to accept both RGB (a red, green and blue colour model) and infrared (sensitive to short-wavelength infrared radiation at around 880nm) images acquired by a range of devices, including a fixed imaging platform using a Nikon D90 digital camera (Fig. 1a), smartphones (e.g. an iPhone, Fig. 1b), or a mobile version CropQuant [32] equipped with either a *Pi* NoIR (no infrared filter) sensor or an RGB sensor (Fig. 1c). When taking pictures, users need to ensure that the camera covers the regions of interest (ROI), i.e. a whole tray (Fig. 1d) or a pot region (Fig. 1e). Red circular stickers (4mm in radius in our case) shall be applied to the four corners of a tray or a pot (Fig. 1b). In doing so, Leaf-GP can extract ROI from a given raw image and then convert measurements from pixels to metric units (i.e. millimetre, mm). Both raw and processed image data can be loaded and saved by Leaf-GP on personal computers (PCs), HPC, or cloud-based computing storage (Figs. 1f&g).

**Figure 1.**
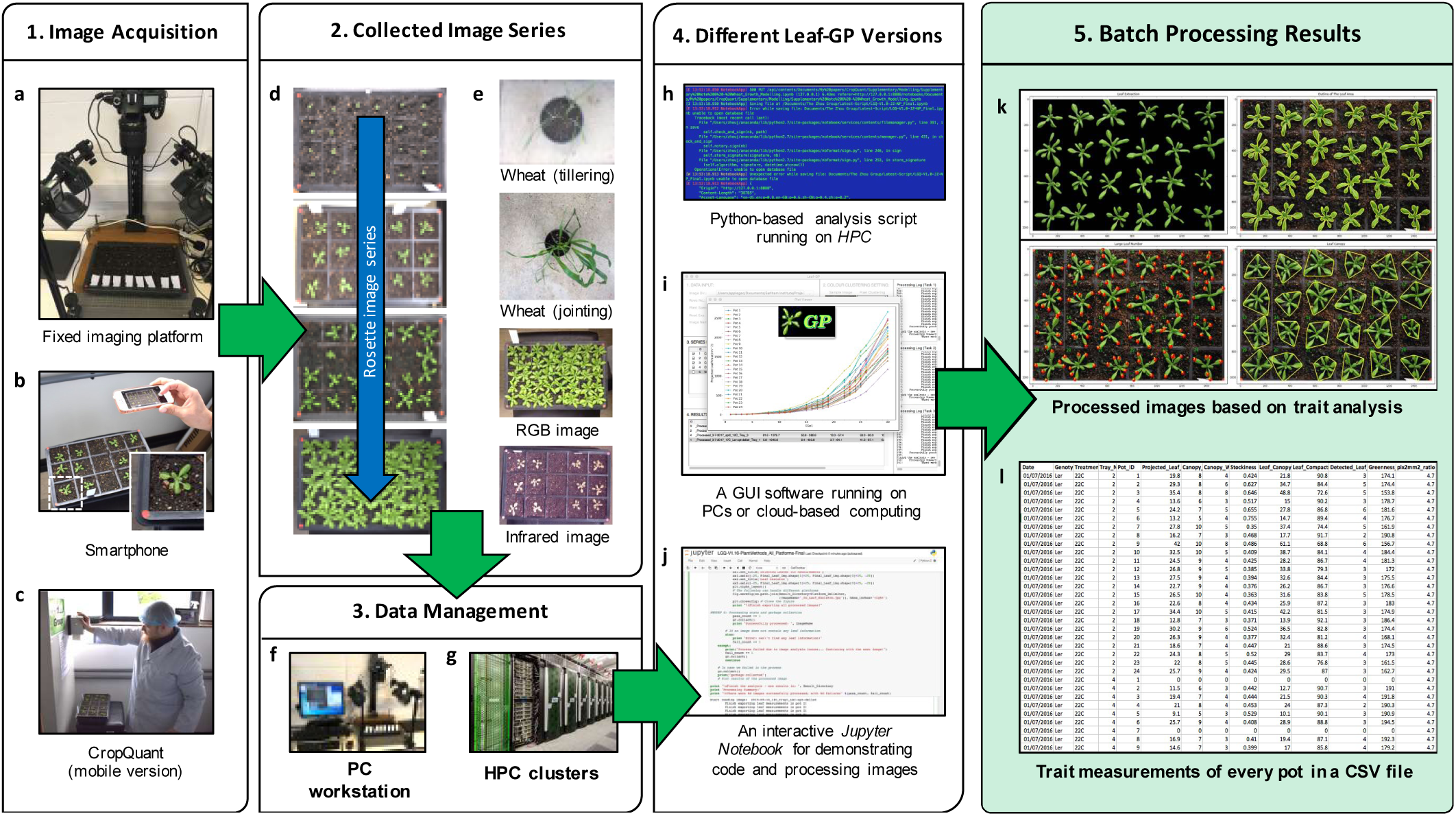
An overview of how to utilise Leaf-GP in plant growth research. (a-c) A range of imaging devices, including a fixed imaging platform, smartphones, or a mobile version CropQuant equipped with either a *Pi* NoIR sensor or an RGB sensor. (d-e) The regions of a tray or a pot need to be covered. (f-g) Both raw and processed image data can be loaded and saved by Leaf-GP on PCs, HPC clusters, or cloud-based computing storage. (h-j) Three versions of Leaf-GP, including HPC, GUI and a *Jupyter Notebook*. (k-l) Processed images highlighting key growth phenotypes and CSV files containing trait measurements are produced after the batch image processing.

As different research groups may have access to dissimilar computing infrastructures, we developed three versions of Leaf-GP to enhance the accessibility of the software: (1) for users utilising HPC clusters, a Python-based script was developed to perform high-throughput trait analysis through a command-line interface (Fig. 1h), which requires relevant scientific and numeric libraries such as SciPy [33], computer vision (i.e. the Scikit-image library [34] and the OpenCV library [35]), and machine learning libraries (i.e. the Scikit-learn library [36]) pre-installed on the clusters; (2) for users working on desktop PCs, a GUI-based (graphic user interface) software application was developed to incorporate batch image processing, multiple trait analysis, and results visualisation in a user-friendly window (Fig. 1i); and, (3) for computational biologists and computer scientists who are willing to exploit our source code, we created an interactive *Jupyter Notebook* (Fig. 1j, also known as the iPython Notebook, see Additional File 1) to explain our multilevel trait analysis workflow and how to modulate code to improve algorithm readability. In particular, we have enable the *Notebook* version to process large image series via a *Jupyter* server, which means users can carry out batch image processing directly using the Notebook version. Due to software distribution licensing issues, we recommend users to install the Anaconda Python distribution (Python 2.7 version) and OpenCV (v2.4.11) libraries before using Leaf-GP. Application File 2 explains the step-by-step procedure of how to install Python and necessary libraries for our software.

After trait analysis, two types of output results are generated. First, *processed images* (Fig. 1k), which includes pre-processing results, calibrated images, colour clustering, and figures exhibiting key growth traits such as leaf outlines, leaf skeletons, detected leaves, and leaf canopy (Additional File 3). Second, *a CSV file* (comma-separated values, Fig. 1l), containing image name, experimental data, pot ID, pixel-to-mm ratio, and biologically relevant outputs including projected leaf area (mm^2^), leaf perimeter, canopy length and width (in mm), stockiness (%), leaf canopy size (mm^2^), leaf compactness (%), the number of leaves, and vegetative greenness (Additional File 4).

### The GUI of Leaf-GP

As plant researchers commonly use PCs for their analyses, we develop the Leaf-GP GUI version based on Python’s native GUI package, Tkinter [37]. The version can operate on different platforms (e.g. Windows and Mac OS) and the default resolution of the main window is set to 1024x768 pixels, so that it can be compatible with earlier operating systems (OS) such as Windows 7. Figure 2 illustrates how to utilise the GUI window to process multiple growth image series (five series were imported with four processed). A high-level analysis workflow of Leaf-GP is presented in Figure 2a, containing five steps: (1) data selection, (2) image pre-processing, (3) global ROI segmentation (i.e. at image level), (4) local trait analysis (i.e. at the pot level), and (5) results output. To explain functions and procedures developed for the workflow, we also prepared a detailed UML (unified modelling language) activity diagram [38] that elucidates stepwise actions, which includes software engineering activities such as choice, iteration, and concurrency to enable the batch processing of large image datasets (Additional File 5).

**Figure 2.**
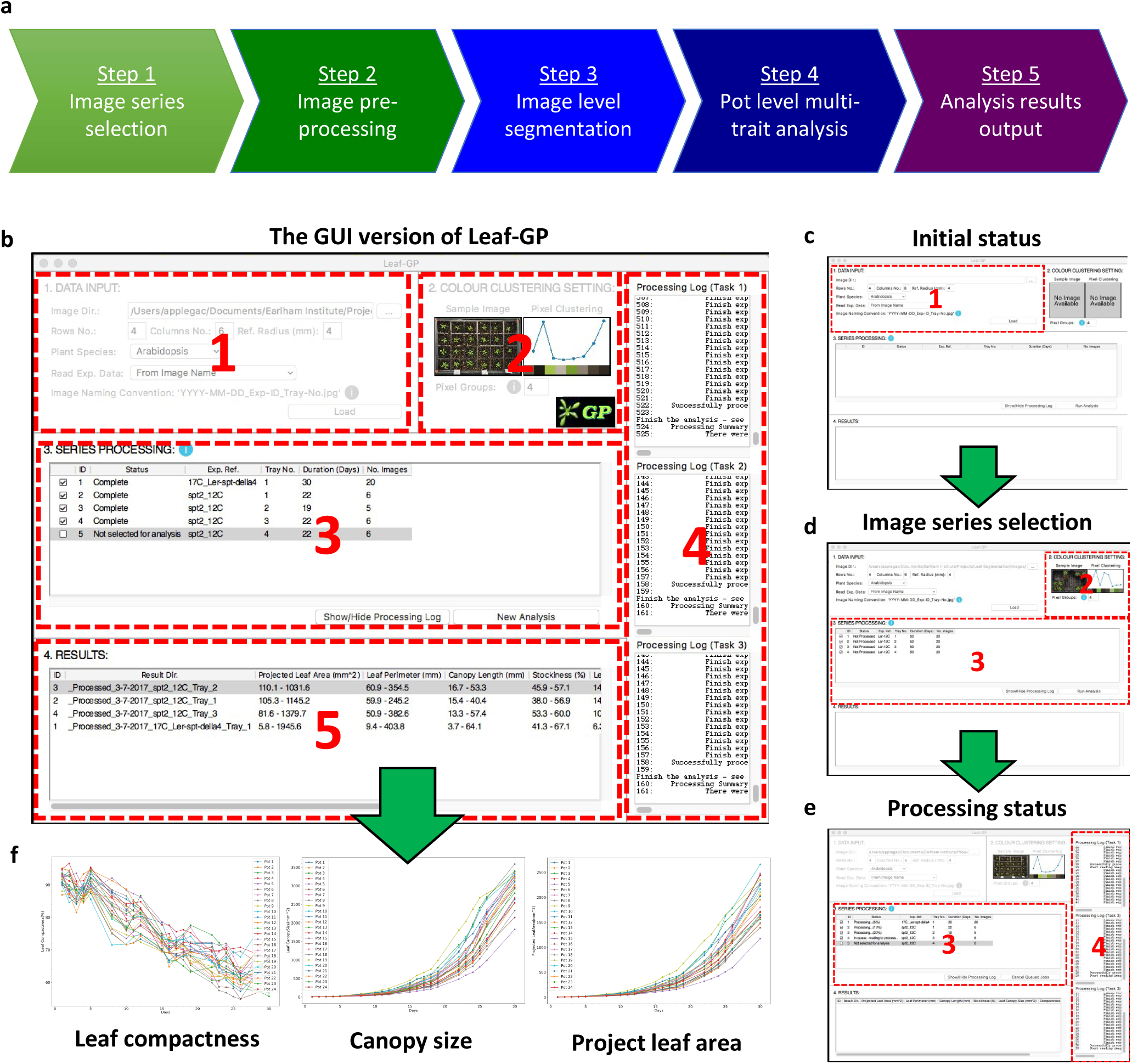
The analysis workflow and the GUI of Leaf-GP. (a) The high-level analysis workflow of Leaf-GP contains five main steps. (b) Five self-explanatory sections designed to integrate the analysis workflow into the GUI version of Leaf-GP. (c) The initial status of the GUI. (d) The screenshot after selecting image series. (e) The screenshot when image datasets are being processed in parallel. (f) Growth-related trait plots can be generated by clicking the associated cell in the Results table.

Figure 2b shows five self-explanatory sections designed to integrate the above analysis steps into the GUI version of the software, including: Data Input, Colour Clustering Setting, Series Processing, Processing Log (a hidden section), and Results Section. To analyse one or multiple image series, users just need to follow these sections sequentially. Also, a number of information icons (coloured blue) have been included to explain how to enter input parameters. Figure 2b demonstrates a screenshot of Leaf-GP after it has finished processing four image series.

#### Section 1 – Data Input

To simplify the data input phase, we only require users to enter essential information regarding their images and associated experiments. To complete the section (Fig. 2c), the user first needs to choose a directory (“Image Dir.”) which contains captured image series. Then, the user shall enter parameters in the “Row No.” and “Column No.” input boxes to define the layout of the tray used in the experiment as well as “Ref. Radius (mm)” to specify the radius of the red stickers. Finally, the user needs to select from “Plant Species” and “Read Exp. Data” dropdowns. All inputs will be verified upon entry to ensure only valid parameters can be submitted to the core algorithm.

In particular, the “Read Exp. Data” dropdown determines how Leaf-GP reads experiment metadata such as imaging date, treatments and genotypes. For example, choosing the “From Image Name” option allows the software to read information from the filename, selecting the “From Folder Name” option will extract metadata from the directory name, whereas the “No Metadata Available” selection will group all images as an arbitrary series for trait analysis. This option allows users to analyse images that are *not* following any data annotation protocols. Although not compulsory, we developed a simple naming convention protocol (Additional File 6) to assist users to annotate image names or folder names tailored for Leaf-GP.

#### Section 2 – Colour Clustering Setting

Once the data input phase is completed, the user can click the ‘Load’ button to initiate series sorting, which will populate the *Colour Clustering Setting* section automatically (Fig. 2d). A sample image from the midpoint of a given series will be chosen by the software, i.e. the image represents the colour groups in the middle of the plant growth. The image is then downsized and processed by a simple k-means method [36], producing a clustering plot and a *k* value that populates in the “Pixel Groups” input box. The user can override the *k* value in the “Pixel Groups” input box; however, to reduce the computational complexity, Leaf-GP only accepts a maximum value of 10 (i.e. 10 representative colour groups) and a minimum value of 3 (i.e. three colour groups) when conducting trait analysis. The generated *k* value (between 3 and 10) will be passed to the core analysis algorithm when the batch processing starts.

#### Sections 3&4 – Series Processing

In the *Series Processing* section (Fig. 2e), the software fills the processing table with information that can help users identify different experiments, including experiment reference (“Exp. Ref.”), the tray number (“Tray No.”), and the number of images in a series (“No. Images”). To improve the appearance of the table, each column is resizable. Checkboxes are prepended to each recognised series. Users can toggle one or multiple checkboxes to specify how many experiments will be processed. If the ‘No Metadata Available’ option is selected (see the *Data Input* section), information such as “Exp. Ref.” and “Tray No.” will not be populated.

The initial status of each processing task (“Status”) is *Not Processed*, which will be updated constantly during the image analysis. When more than one experiment is selected, Python’s thread pool executor function will be applied, so that these experiments can be analysed simultaneously in multiple cores in the central processing unit (CPU). We have limited up to *three* analysis threads (see the right of Fig. 2e), because many Intel processors comprise *four* physical cores and conducting parallel computing can have a high demand of computing resources (e.g. storage, CPU and memory), particularly during the batch processing when raw image datasets are big.

Once the processing table is filled, the user can click the ‘Run Analysis’ button to commence the analysis. Figure 2b shows the screenshot when five experiments (i.e. five image series) are recognised and four of them are analysed. Due to the multi-task design of Leaf-GP, we only allowed three series running in parallel. Throughout the analysis, the ‘Status’ column will be continually updated, indicating how many images have been processed. It is important to note that, although Leaf-GP was designed for parallel computing, some functions used in the core algorithm are not thread-safe, indicating they can only be executed by one thread at a time. Because of this limit, we have utilised lock synchronisation mechanisms to protect code blocks (i.e. procedures or functions), so that these thread-unsafe blocks can only be executed by one thread at a time. In addition to the processing status, more analysis information can be viewed by opening the *Processing Log* section (to the right of Fig. 2e), which can be displayed or hidden by clicking the ‘Show/Hide Processing Log’ button on the main window.

#### Section 5 – Results

When all processing tasks are completed, summary information will be appended to the Results section, including processing ID and a link to the result folder which contains the CSV file and all processed images (“Result Dir.”). Depending on which species (i.e. *Arabidopsis* rosette or wheat) is selected, trait plots will be generated to show key growth phenotypes (e.g. the projected leaf area, leaf perimeter, leaf canopy size, leaf compactness, and leaf numbers) by clicking on the associated cell in the Results table (Fig. 2f). The range of phenotype measurements is also listed in the *Results* section. The GUI version also saves processing statistics, for example, how many images have been successfully analysed and how many images have been declined, together with related error or warning messages in a log file for debugging purposes.

### Core trait analysis algorithms

Multiple trait analysis of *Arabidopsis* rosettes and wheat plants is the core part of Leaf-GP. Not only does it utilise advance computer vision algorithms for automated analysis, it also encapsulates feature extraction and phenotypic analysis methods that are biologically relevant to growth phenotypes. In the following sections, we explain the core analysis algorithm in detail.

#### Step 2 – Pre-processing and calibration

Different imaging devices, camera positions and even lighting conditions can cause quality variance during image acquisition. Hence, it is important to calibrate images before conducting automated trait analysis. We developed a pre-processing and calibration procedure as shown in Figure 3. We first resized each image (Fig. 3a) to a fixed resolution so that the height (i.e. y-axis) of all images in a given series could be fixed. A `rescale` function in Scikit-image was used to dynamically transform the image height to 1024 pixels (Fig. 3b). After that, we created a **`RefPoints`** function (*Function_2* in Additional File 1) to detect red circular markers attached to the corners of a tray or a pot region. To extract these markers robustly under different illumination conditions, we designed *g(x, y)*, a multi-resholding function to segment red objects derived from a single-colour extraction approach [39]. The function defines which pixels shall be retained (intensity is set to 1) and which pixels shall be discarded (intensity is set to 0) after the thresholding:

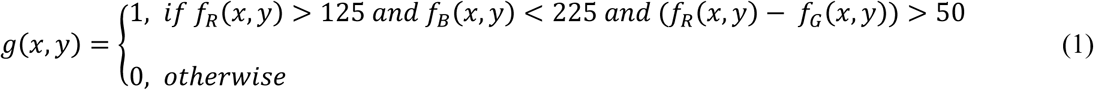

where *f*_*R*_*(x, y)* is the red channel of a colour image, *f*_*B*_*(x, y)* represents the blue channel and *f*_*G*_ *(x, y)* the green channel. The result of the function is saved in a reference binary mask.

**Figure 3.**
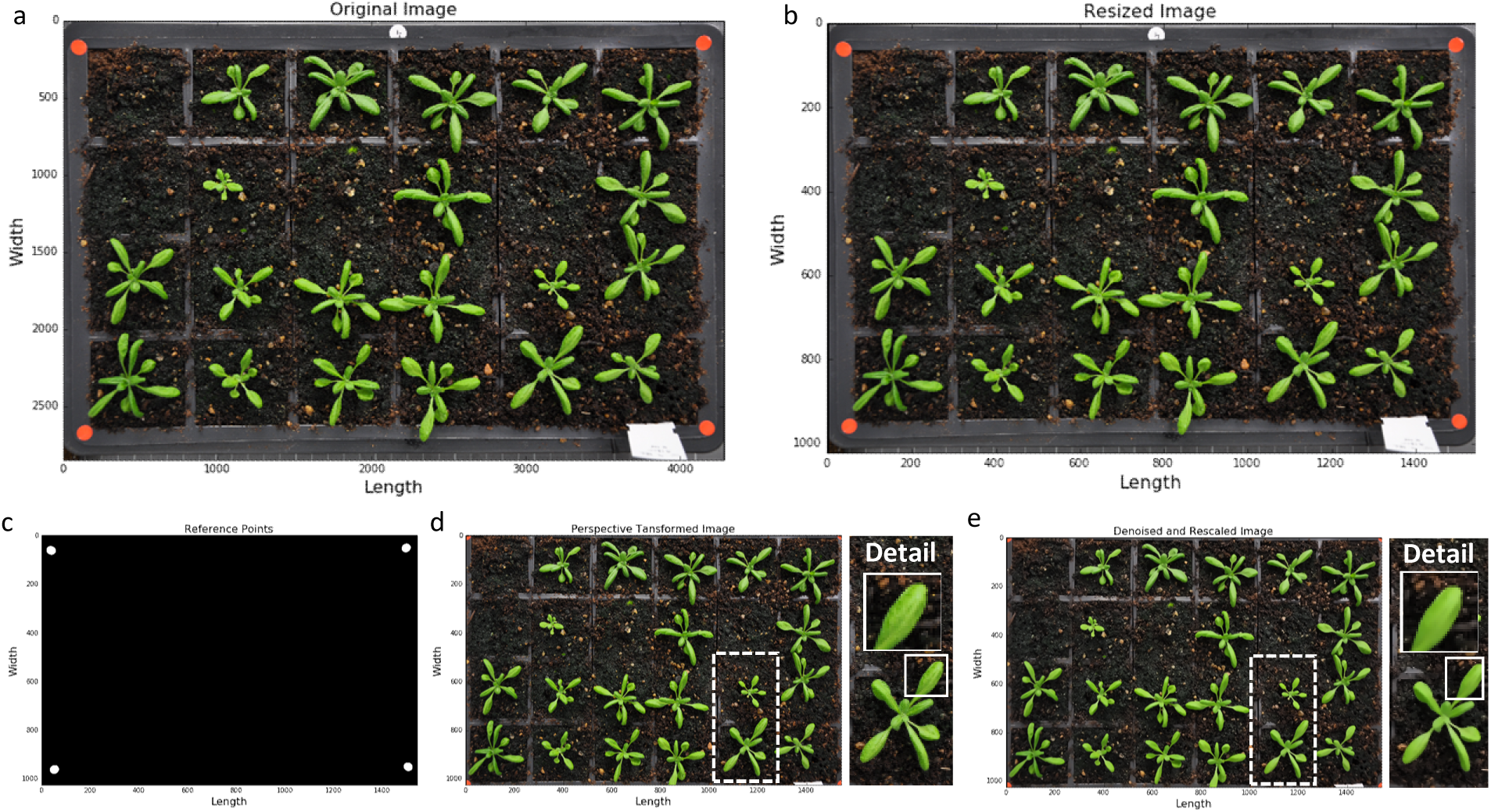
The step of image pre-processing and calibration. (a-b) Fix the height (i.e. y-axis) of all images in a given series. (c) Detect red circular markers. (d) Extract ROI from the original image. (e) Denoise the image to smooth leaf surface for the global leaf segmentation.

We then used the **`regionprops`** function in Scikit-image to measure morphological features of the reference-point mask to filter out false positive items. For example, if an object’s area, eccentricity or solidity readings do *not* fit into the characteristics of a circle, this object will be discarded. After this step, only genuine circular objects are retained (Fig. 3c) and the average radius (in pixels) of these circular objects is converted to mm units (the radius of the red markers is 4mm). To extract the tray region consistently, we developed a tailored algorithm called **`PerspectiveTrans_2D`** (*Function_5* in Additional File 1), using **`getPerspectiveTransform`** and **`warpPerspective`** functions in OpenCV to retain the region that is enclosed by the red markers (Fig. 3d). Finally, we employed a non-local means denoising function called **`fastNlMeansDenoisingColored`** in OpenCV to smooth leaf surface for the following global leaf ROI segmentation (Fig. 3e).

#### Step 3 – Global leaf ROI segmentation

Besides imaging related issues, changeable experimental settings could also cause issues for automated trait analysis. Figures 4a-d illustrate a number of problems we have encountered whilst developing Leaf-GP. For example, the colour and texture of the soil surface can change considerably between different experiments, especially when gritty compost and other soil types are used (Figs. 4a&b); sometimes plants are *not* positioned in the centre of a pot (Fig. 4b), indicating leaves that cross over to adjacent pots should be segmented; algae growing on the soil has caused false detection due to their bright green colour (Figs. 4c&d); finally, destructive harvest for weighing biomass could occur from time to time throughout an experiment, indicating the core analysis algorithm needs to handle random pot disruption robustly (Fig. 4d). To address the above technical challenges, we developed a number of computer vision and simple machine-learning algorithms based on open scientific libraries. Results of our software solutions integrated in Leaf-GP can be seen to the right of Figures 4a-d.

**Figure 4.**
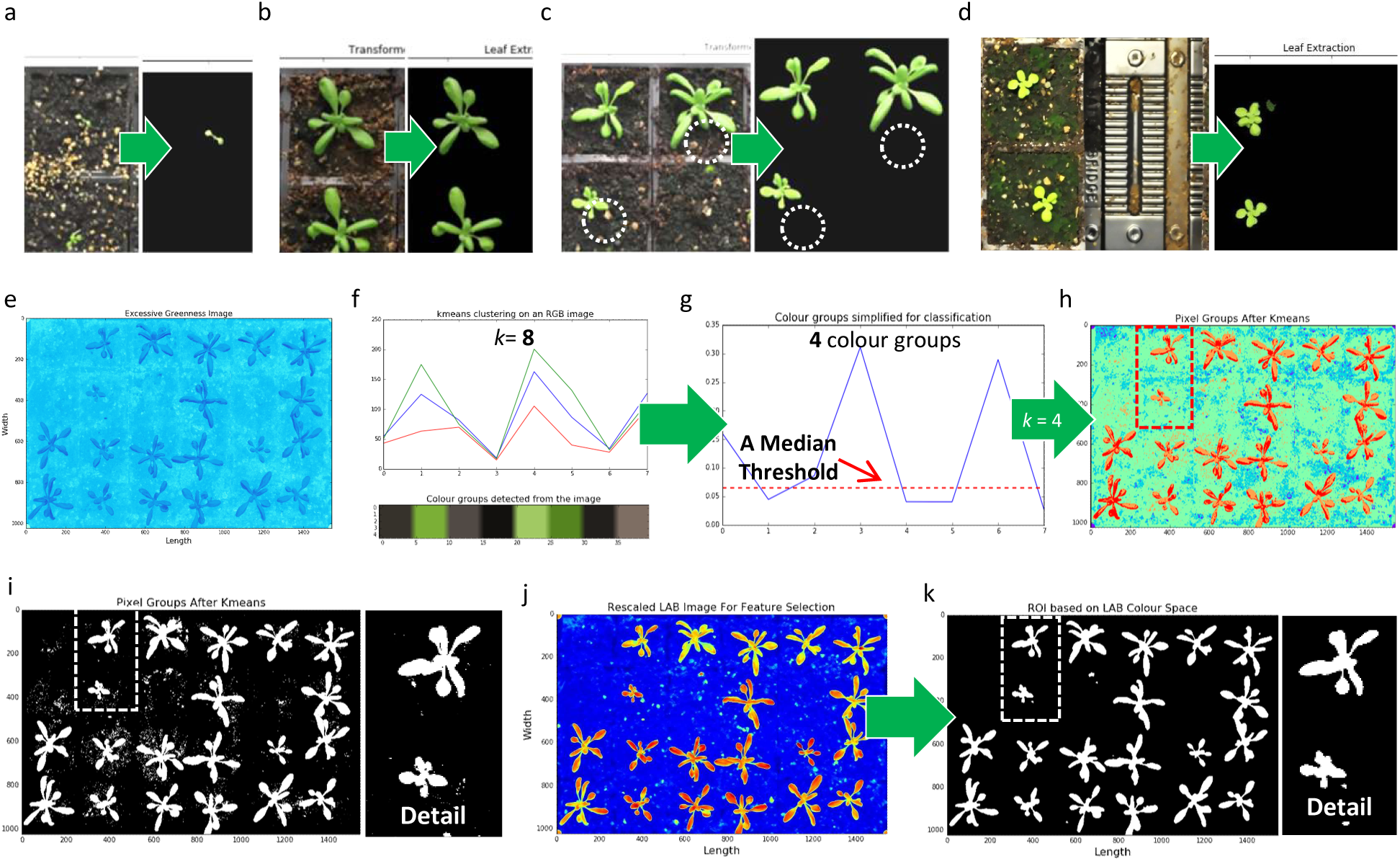
The step of defining global leaf ROI. (a-d) A number of experiment-related problems encountered whilst developing Leaf-GP (to the left of the figures) and results of our solutions (to the right of figures). (e) A pseudo vegetative greenness image. (f-g) Using KMeans to estimate how many colour groups can be classified from a given colour image. (h) The classification result of the KMeans approach based on the pseudo vegetative greenness picture, highlighting greenness values in red pixels. (i) A global adaptive Otsu thresholding used to generate a global leaf ROI binary mask. (j-k) Lab colour space used to extract leaf ROI objects at the image level.

The first approach we developed is to establish a consistent approach to extract pixels containing high values of greenness (i.e. leaf regions) from an RGB image robustly. Using a calibrated image, we computed vegetative greenness *G*_*V*_*(x, y)* [13] based on excessive greenness *Ex*_*G*_ *(x, y)* and excessive red *Ex*_*R*_*(x, y)* indices. The pseudo vegetative greenness image (*G*_*V*_, Figure 4e) is produced by equation 2, based on which we implemented a **`compute_greenness_img`** function (*Function_8* in Additional File 1) to transfer an RGB image into a *G*_*V*_ picture. Excessive greenness is defined by equation 3 and excessive red is defined by equation 4:

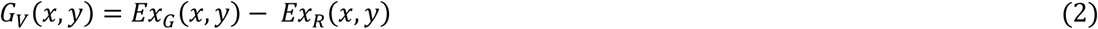

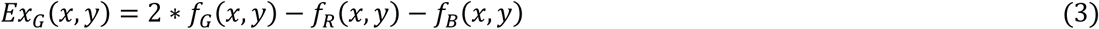

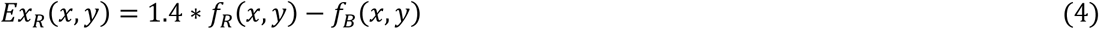

where *f*_*R*_*(x, y)* is the red channel of a colour image, *f*_*B*_*(x, y)* represents the blue channel, and *f*_*G*_ *(x, y)* the green channel.

After that, we applied a simple unsupervised machine learning algorithm called **`KMeans`** (default *k* = 8 was used, assuming 8 representative colour groups in a given image) and **`KMeans.fit`** in Scikit-learn to estimate how many colour groups can be classified (Fig. 4f, *Function_8.1* in Additional File 1). We chose a median threshold (red dotted line) to segment the colour clustering result and obtained the *k* value to represent the number of colour groups (Fig. 4g). Also, this process has been integrated into the GUI version (i.e. the *Colour Clustering Setting* section). Utilising the *k* value (e.g. *k* = 4, Fig. 4g), we designed a **`kmeans_cluster`** function (*Function_9* in Additional File 1) to classify the pseudo vegetative greenness picture, highlighting greenness values in red pixels (Fig. 4h). A global adaptive Otsu thresholding [40] was used to generate an image level leaf ROI binary mask (Fig. 4i). However, the simple machine learning approach could produce miss-detected objects due to complicated colour presentations during the plant growth period (e.g. Figs. 4a-d). For example, the k-means approach performed well when the size of the plants is between 25-75% of the size of a pot, but created many false detections when leaves are tiny or the background is complicated. Hence, we designed another approach to improve the detection based on the k-means approach.

We employed Lab colour space [41], which incorporates lightness and green-red colour opponents to refine the detection. We created an internal procedure called **`LAB_Img_Segmentation`** (*Function_7* in Additional File 1) to transfer RGB images into Lab images using the **`color.rgb2lab`** function in Scikit-image, based on which green pixels were featured in a non-linear fashion (Fig. 4j). Again, a global adaptive Otsu thresholding was applied to extract leaf objects and then a Lab-based leaf region mask (Fig. 4k). Finally, we combined the Lab-based mask with the k-means mask as the output of the phase of global leaf ROI segmentation.

#### Step 4.1 – Pot level segmentation

To measure growth phenotypes in a given pot over time, plants within each pot need to be monitored over time. Using the calibrated images, we have defined the tray region, based on which we constructed the pot framework in the tray. To accomplish this task, we designed an iterative layout drawing method called **`PotSegmentation`** (*Function_5* in Additional File 1) to generate anti-aliased lines using the **`line_aa`** function in Scikit-image to define the pot layout (Fig. 5a).

**Figure 5.**
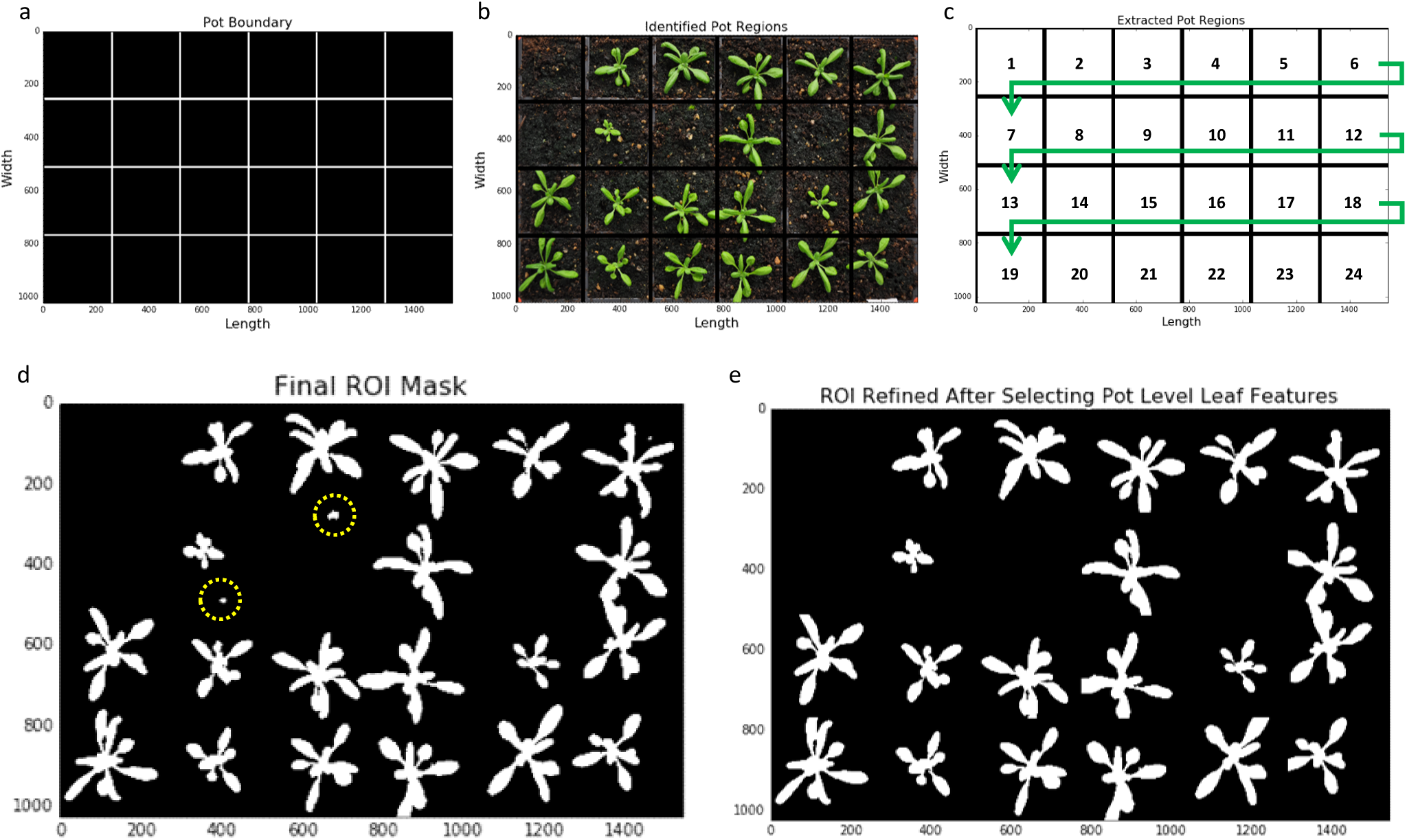
The step of conducting pot level segmentation in a sequential manner. (a) Depending on the number of rows and columns, generate anti-aliased lines to define the pot layout. (b) Segmented a given image into a number of sub-images. (c) The sequence to go through each pot an in an iterative approach. (d-e) Apply a local detection method to improve the result of leaf detection.

After constructing the framework, we segmented the leaf growth image into a number of sub-images (Fig. 5b), so that plant can be analysed locally, at the pot level. We developed an iterative analysis approach to go through each pot with the sequence presented in Figure 5c. Within each pot, we conducted a local leaf detection method. For example, although combining leaf masks produced by the machine learning (Fig. 4i) and the Lab colour space (Fig. 4k) approaches, some false positive objects may still remain (Fig. 5d). The local leaf detection can enable us to employ pot-level contrast and intensity distribution [42], weighted image moments [43], texture descriptor [44], and leaf positional information to examine each sub-image to refine the leaf detection (Fig. 5e). This local feature selection method (detailed in the following sections) can also help us decrease the computational complexity (i.e. memory and computing time), as analysis is carried out within smaller sub-images.

#### Step 4.2 – Local multiple trait measurements

Utilising the refined local leaf masks at the pot level (Fig. 6a), a number of growth phenotypes could be quantified reliably. Some of them are enumerated briefly as follows:

**Figure 6.**
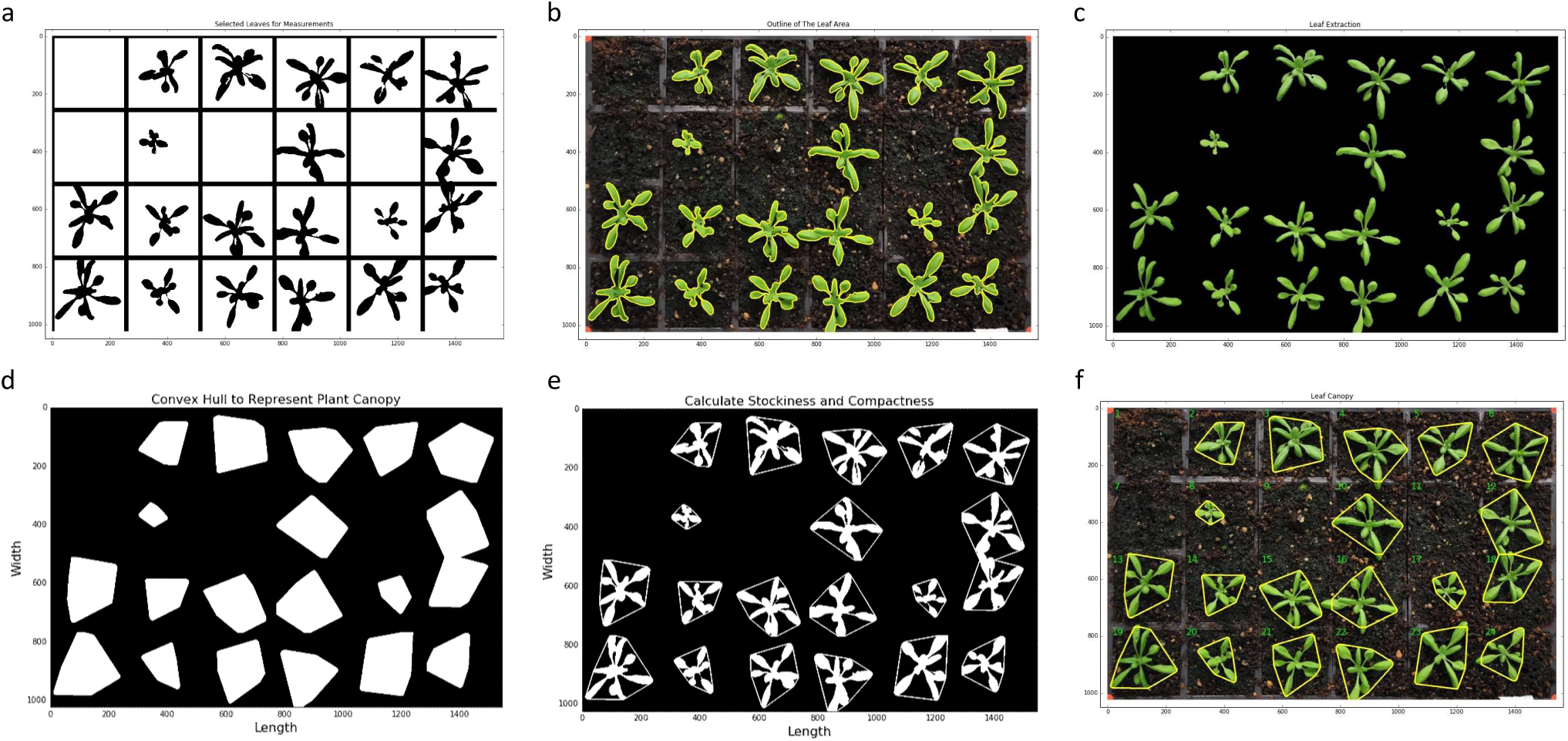
The step of measuring multiple growth traits. (a) Refined leaf masks for every pot. (b) Contours generated to outline the leaf region. (c) Green pixels enclosed by the contours are totalled for computing the size of the projected leaf area. (d) Convex hulls created in every pot for calculating leaf canopy. (e) Stockiness calculated based on the ratio between the plant projected area and the leaf perimeter. (f) Leaf Compactness computed based on the ratio between the projected leaf area and the area of the convex hull.

1. “Projected Leaf Area (mm^2^)” measures the area of an overhead projection of the plant in a pot. While implementing the function, the **`find_contours`** function in Scikit-image is used to outline the leaf region (coloured yellow in Fig. 6b). Green pixels enclosed by the yellow contours are totalled to compute the size of the projected leaf area (Fig. 6c). Pixel-based quantification is then converted to mm units based on the pixel-to-mm exchange rate. This trait is a very reliable approximation of the three-dimensional (3D) leaf area and has been used in many growth studies [20,22,45].
2. “Leaf Perimeter (mm)” is calculated based on the length of the yellow contour line that encloses the detected leaf region. Again, pixel-based measurements are converted to mm units, which are then used to compute the size change of a plant over time.
3. “Daily Relative Growth Rate (%)” (Daily RGR) quantifies the speed of plant growth. Derived from the RGR trait described previously [19,46], the Daily RGR here is defined by equation 5:

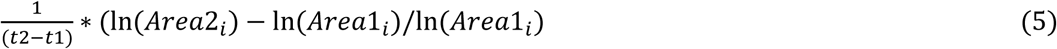

where *ln* is natural logarithm, *Area*1_*i*_ is the projected leaf area in pot *i* in the previous image, *Area*2_*i*_ is the leaf area in pot *i* in the current image, and *(t*2 - *t*1) is the duration (in days) between the two consecutive images.
4. “Leaf Canopy (mm^2^)” expresses the plant canopy region that is enclosed by a 2D convex hull in a pot [19,20,22]. The convex hull was generated using the **`convex_hull_image`** function in Scikit-image, enveloping all pixels that belong to the plant with a convex polygon [47]. Figure 6d presents all convex hulls created in a given tray. As described previously [19], this trait can be used to define the coverage of the leaf canopy region as well as how the petiole length changes during the growth period.
5. “Stockiness (%)” is calculated based on the ratio between the plant projected area and the leaf perimeter (Fig. 6e). It is defined as *(4*π * *Area*_*i*_)/(2π * *R*_*i*_)^2^, where *Area*_*i*_ is the projected leaf area detected in pot *i* and *R*_*i*_ is the longest radius (i.e. major axis divided by 2) of the convex hull polygon in pot *i*. This trait (0-100%) has been used to measure how serrated a plant is, which can also indicate the circularity of the leaf region (e.g. a perfect circle will score 100%).
6. “Leaf Compactness (%)” is computed based on the ratio between the projected leaf area and the area of the convex hull enclosing the plant [20,22]. Figure 6f shows how green leaves are enclosed by yellow convex hull outlines that calculates the leaf compactness trait.
7. “Greenness” monitors the normalised greenness value (0-255) of the leaf canopy, i.e. the convex hull region. A rescaled Lab image is used to provide the greenness reading, so that we could minimise the background noise caused by algae and soil types. Greenness can be used to study plant growth stages such as vegetation and flowering.

#### Step 4.3 – Leaf number detection

As the number of rosette leaves is popularly used to determine key growth stages for *Arabidopsis* [15], we therefore designed a leaf structure detection algorithm to provide a consistent reading of traits such as the number of detected leaves and the number of detected long or large leaves over time. This algorithm comprises of a 2D topological skeletonisation algorithm (*Function_10* in Additional File 1) and a leaf outline sweeping method (*Function_11* in Additional File 1).

Figure 7a demonstrates the result of the skeletonisation approach, which utilises the **`skeletonize`** function in Scikit-image to extract 2D skeletons from the leaf masks in each pot. The skeletons can be used to quantify the structural characteristics of a plant, including the number of leaf tips and branching points of a plant. For example, we implemented a **`find_end_points`** function to detect each leaf tip (i.e. end point) in a plant skeleton using the **`binary_hit_or_miss`** function in the SciPy library to match the four possible 2D matrix representations:

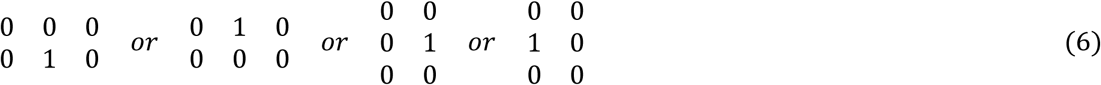

**Figure 7.**
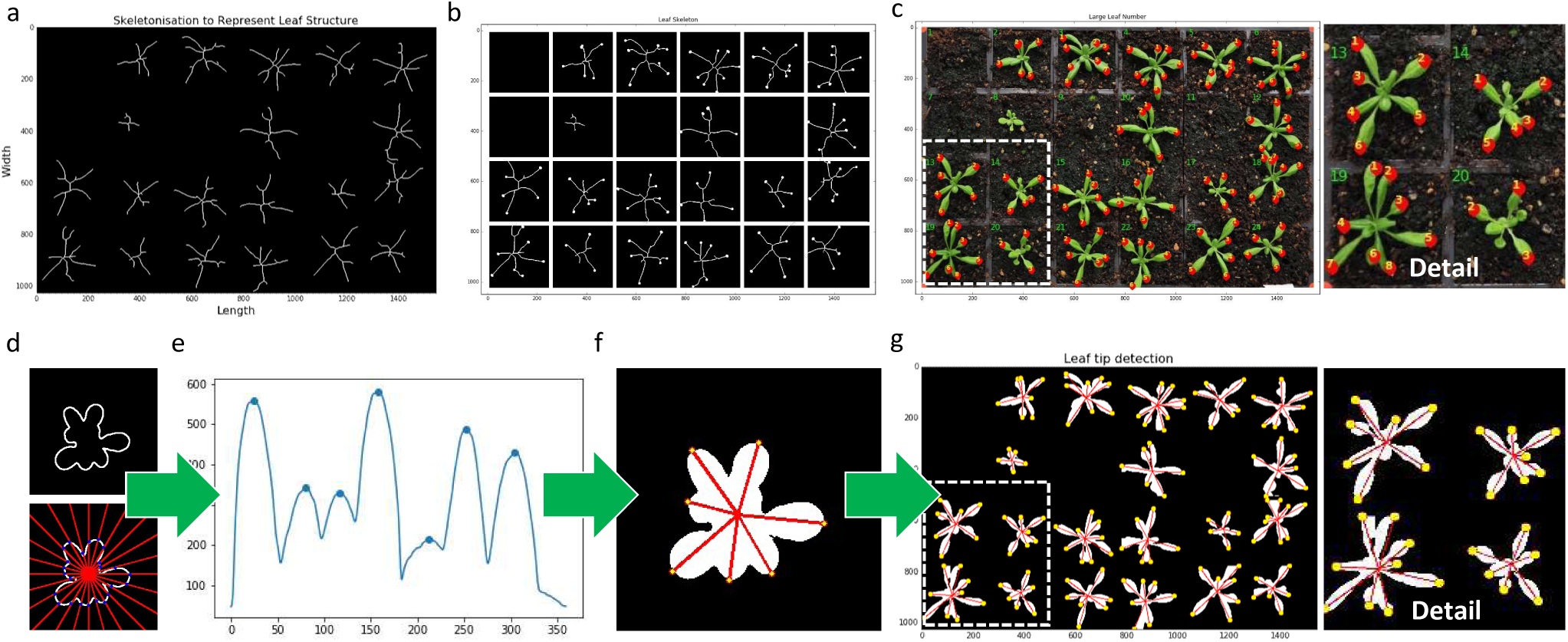
The step of detecting leaf structure. (a) The result of a 2D skeletonisation approach to extract leaf structure. (b) Detect end points of the leaf structure which correlates with leaf tips. (c) Large or long rosette leaves identified if they are over 50% or 70% of the final size. (d-e) Generate a distance series to represent the distance between the plant centroid and its leaf contour, at angles between 0 and 359 degrees with a 15-degree interval. (f-g) The number of detected peaks are used to represent the number of leaf tips.

The **`find_end_points`** function outputs 2D coordinates of end points that correlates with leaf tips (Fig. 7b). Furthermore, the function can be employed for novel trait measurements, for instance, large or long rosette leaves can be identified if they are over 50% or 70% of the final size (Fig. 7c and *Step_4.4.2.7* in Additional File 1). To accomplish this, we evaluated the leaf skeleton as a weighted graph and then treated: (1) the skeleton centroid and end points as *vertices* (i.e. *nodes*), (2) lines between the centre point and end points as *edges*, and (3) the leaf area and the length between vertices as *weights* assigned to each *edge*. Depending on the experiment, if the *weights* are greater than a predefined threshold (i.e. over 15mm in length and 100mm^2^ in leaf size in our case), the associated leaf will be recognised as a long or large leaf.

As the skeletonisation approach could miss some small leaves if they are close to the plant centroid or partially overlapping with other leaves, we implemented a **`leaf_outline_sweeping`** procedure to establish another approach to detect the total leaf number based on the distance between the plant centroid and any detected leaf tips. This procedure is based on a published leaf tip identification algorithm [5]. We improved upon the algorithm through utilising the leaf boundary mask (i.e. contour) to reduce the computational complexity. For a given plant, the algorithm generates a distance series that represents the squared Euclidean distances from the plant centroid to its contour, at angles between 0 and 359 degrees with a 1-degree interval (for presentation purposes, we only used 15 degree intervals in Fig. 7d). To reduce noise, the algorithm smooths the distance series using a Gaussian kernel (Fig. 7e). A peak detection algorithm called **`PeakDetect`** [48] is integrated in our core analysis algorithm to detect peaks on the distance series (*Step_4.4.2.8* in Additional File 1). The procedure implemented here supports our assumption that the number of peaks can be used to largely represent the number of leaf tips (Figs. 8f&g). When quantifying the total number of leaves, results from both skeleton and outline sweeping approaches are combined to produce a viable measurement.

**Figure 8.**
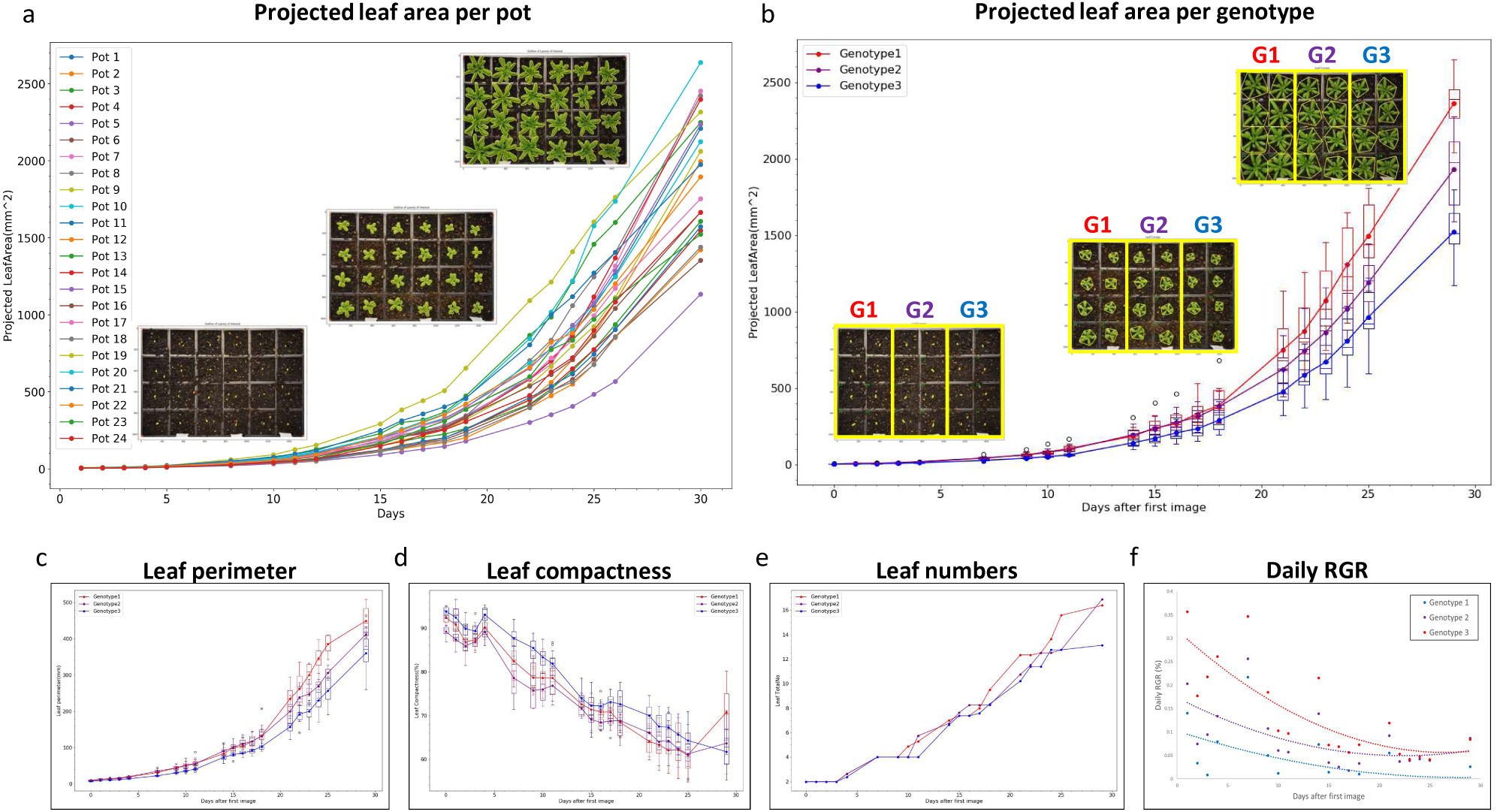
Case study 1: Analysis results of a tray with three genotypes. (a) The projected leaf area trait in 24 pots was quantified by Leaf-GP. (b) The projected leaf area traits divided into three genotype groups. (c-f) A number of growth related traits such as leaf perimeter, compactness, leaf number, and daily RGR of all three genotypes can be statistically differentiated.

## Results

Leaf-GP can facilitate plant growth studies through automating trait analysis and cross-referencing results between experiments. Instead of merely utilising machine learning algorithms to build neural network architecture for pixel clustering or trait estimates [49], we chose an approach that combines simple unsupervised machine learning and advance computer vision algorithms to establish an efficient analysis workflow. This approach has enabled us to select morphological features that are biologically relevant for conducting meaningful ROI segmentation at both image and pot levels. Here, we exhibit three use cases where Leaf-GP were employed to study key growth phenotypes for *Arabidopsis* rosettes and *Paragon* wheat.

### Use case 1 – Tracking three genotypes in a single tray

We applied Leaf-GP to measure growth phenotypes in a tray containing three genotypes L*er* (wildtype), *spt-2*, and *gai-t6 rga-t2 rgl1-1 rgl2-1* (*della4)* at 17°C. Each pot in the tray was monitored and cross-referenced during the experiment. The projected leaf area trait in 24 pots was quantified by Leaf-GP (Fig. 8a) and rosette leaves were measured from stage 1.02 (2 rosette leaves, around 5mm^2^) to stage 5 or 6 (flower production, over 2400mm^2^), a duration of 29 days after the first image was captured.

After dividing the quantification into three genotype groups, we used the projected leaf area readings (Fig. 8b) to verify the previously manually observed growth differences between L*er*, *spt-2*, and *della4* [2,3]. Furthermore, the differences in phenotypic analyses such as leaf perimeter, compactness, leaf number, and daily RGR of all three genotypes can be statistically differentiated (Figs. 8c–f). Particularly for Daily RGR (Fig. 8f), the three genotypes exhibit a wide variety of growth rates that are known to be determined by genetic factors [50]. Based on image series, Leaf-GP can integrate time and treatments (e.g. temperature signalling or chemicals) with dynamic growth phenotypes for cross referencing. We provided the CSV file for *Use Case 1* in Additional File 4, containing trait measurements for each pot over time. The Python script we used to plot and cross-reference either pot- or genotype-based growth phenotypes is provided in Additional File 7, called Leaf-GP plot generator.

### Use case 2 – Two genotypes under different temperatures

We also used our software to detect differences in rosette growth between L*er* (wildtype) and *spt-2* grown at different temperatures, i.e. 12°C and 17°C. Utilising the projected leaf area measurements, we observed that temperatures affect vegetative growth greatly on both lines (Fig. 9a). Similar to previously studied [2,3], lower temperatures can have a greater effect on the growth of *spt-2* than L*er.* Around seven weeks after sowing, the projected leaf area of *spt-2* was around 50% greater on average (1270mm^2^) compared to L*er* (820mm^2^), when grown at 12°C (Fig. 9c). However, when grown in 17 °C, at 36 days-after-sowing *spt-2* had a similar area at around 1200mm^2^, but L*er* had an area of 1000mm^2^, a much smaller difference.

**Figure 9.**
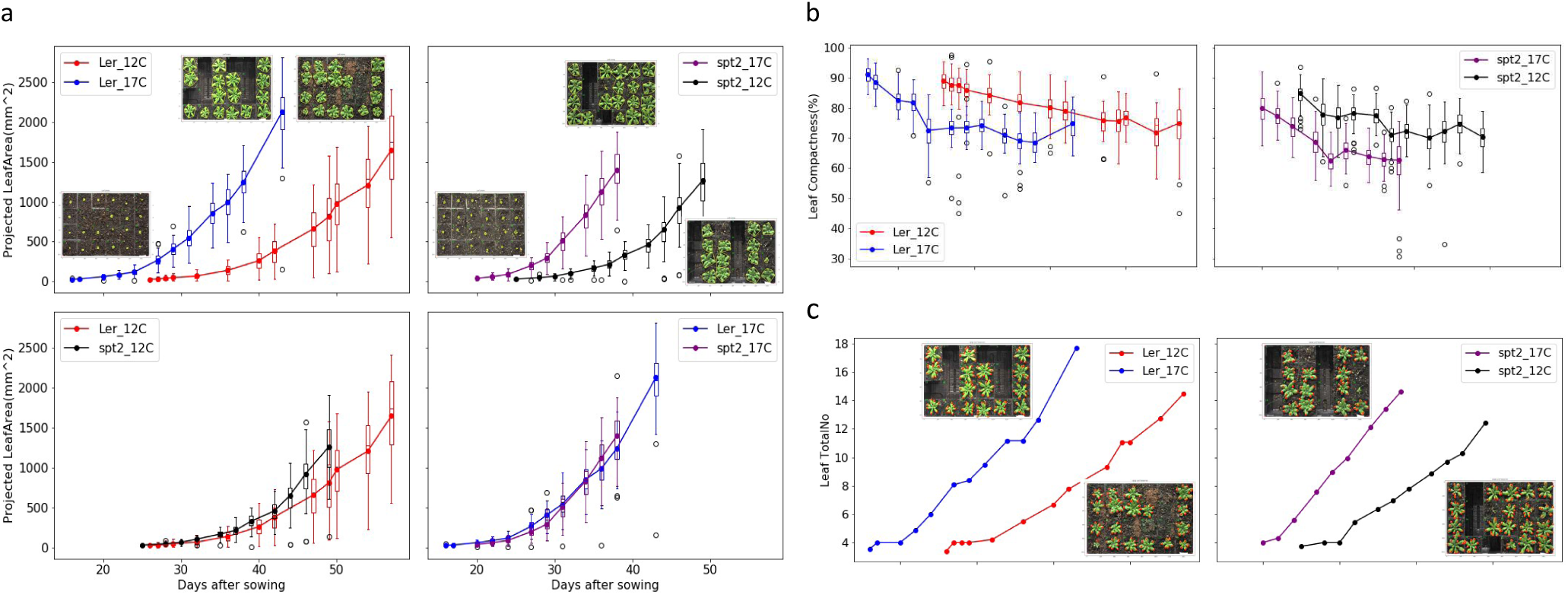
Case Study 2: Analysis results of multiple experiments. (a) The projected leaf area measurements used to observe how temperatures affect vegetative growth on both L*er* and *spt-2*. (b) Plants of both genotypes growing at 12°C had more compact rosettes that those growing at 17°C. *spt-2* was less compact than L*er* in general. (c) The number of leaves produced was greater at the warmer temperature.

As our software can export multiple growth phenotypes, we therefore investigated both linked and independent effects of temperature on wildtype and *spt-2*. For instance, the larger rosette in *spt-2* causes a similar increase in rosette perimeter, canopy length and width, and canopy size. At similar days after sowing, plants of both genotypes grown at 12°C had more compact rosettes that those growing at 17°C (Fig. 9b), and *spt-2* was less compact than L*er* in general. The number of leaves produced was greater at the warmer temperature (Fig. 9c). This ability to easily export a number of key growth traits of interest is useful and relevant to broader plant growth research. We provided detailed processing results (csv files) for the L*er* (12°C and 17°C, Additional File 8) and *spt-2* (12°C and 17°C, Additional File 9) experiments. Results including processed images and CSV files for the two experiments can also be downloaded at https://github.com/Crop-Phenomics-Group/Leaf-GP/releases.

### Use case 3 – Monitoring wheat growth

Another application for which Leaf-GP has been designed is to analyse wheat growth images taken in glasshouses or growth chambers. Similarly, red circular stickers are required to attach to the corners of the pot region so that Leaf-GP can extract ROI and traits can be measured in mm units. Figure 10 demonstrates a proof-of-concept study demonstrating how Leaf-GP has been applied to measure projected leaf area and leaf canopy size based on *Paragon* (a UK spring wheat) image series taken over a 70-day period in greenhouse, from sprouting (Fig. 10b), to tillering (Fig. 10c), and then from booting (Fig. 10e) to heading (Fig. 10f). With a simple and cheap imaging setting, Leaf-GP can precisely quantify key growth phenotypes for wheat under different experimental conditions. Please note that the leaf counting function in Leaf-GP cannot be reliably applied to quantify wheat leaves, because the complicated plant architecture of wheat plants.

**Figure 10.**
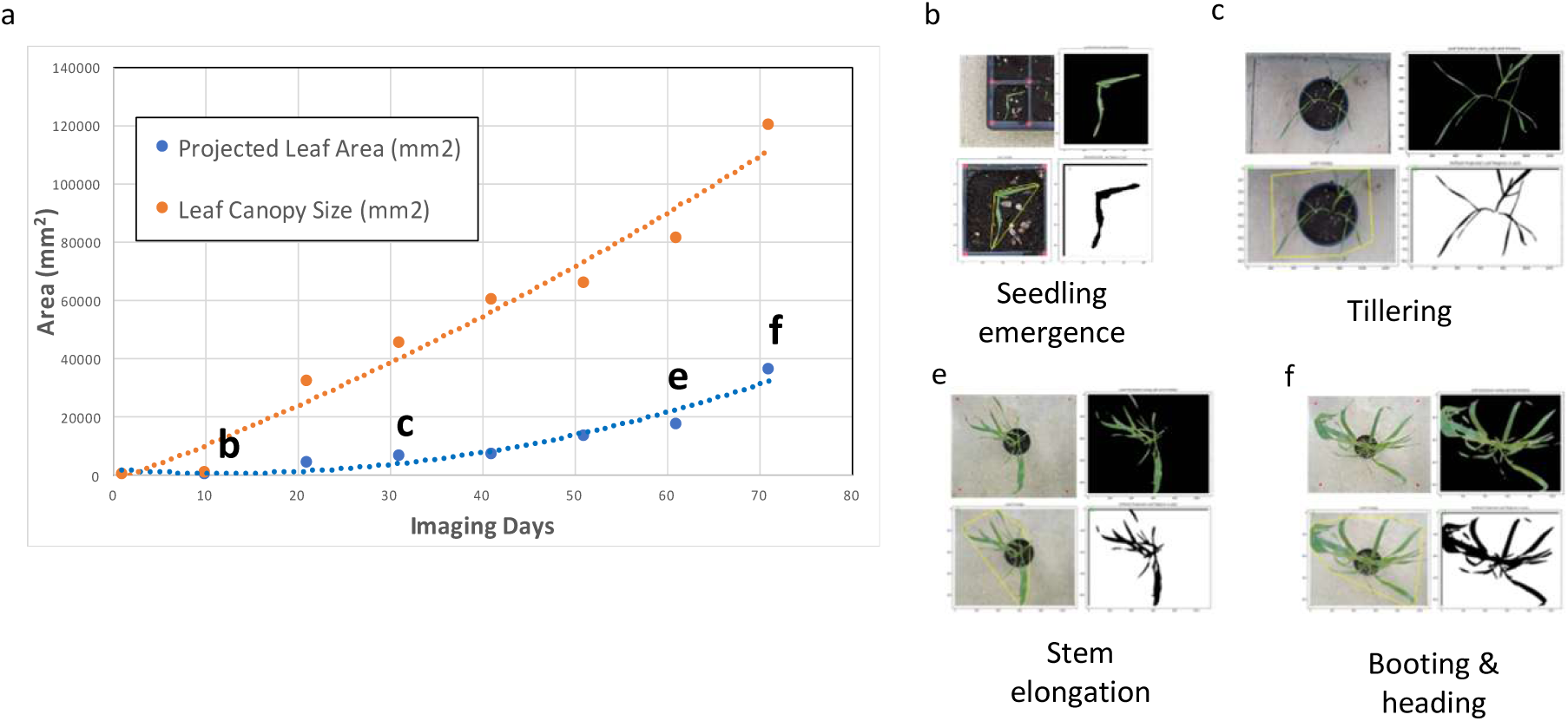
Case Study 3: Applying Leaf-GP on wheat growth studies. (a) A proof-of-concept study of how to measure the projected leaf area and the leaf canopy size based on *Paragon* wheat images, taken over a 70-day period in greenhouse. (b-f) Analysis results generated from sprouting to heading stage.

## Discussion

Different environmental conditions and genetic mutations can impact a plant’s growth and development, making the quantification of growth phenotypes a useful tool to study how plants respond to different biotic and abiotic treatments. Amongst many popularly used growth phenotypes, imaging leaf-related traits is a non-destructive and reproducible approach for plant scientists to record plant growth over time. In comparison with many published image analysis software tools for leaf phenotyping, our software provides a comprehensive solution that is capable of extracting multiple traits automatically from large image datasets; and moreover, it can provide traits analysis that can be used to cross reference different experiments. In order to serve a broader plant research community, we designed three versions of Leaf-GP, including a graphic user interface for PC users, a command-line interface for HPC users, and a *Jupyter Notebook* for computational users. We provide all steps of the algorithm design and software implementation, together with raw and processed datasets we produced for our *Arabidopsis* and *Paragon* wheat studies at NRP.

When developing the software, we particularly considered how to enable different sizes of plant research laboratories to utilise our work for screening large populations of *Arabidopsis* and wheat in response to varied treatments through accessible and low-cost imaging devices. Hence, we paid much attention to software usability (e.g. simple command-line interface or GUI), capability (automatic multiple trait analysis running on different platforms), expandability (open software architecture, new Python-based functions and procedures can be easily added to the software, see the **`PeakDetect`** procedure in Additional File 1), and biological relevance (i.e. the feature extraction approach and processing results are biological relevant). We trust our software is suitable for studying the growth performance of a large number of plant genotypes and treatments with very limited imaging hardware and software resource requirements.

The software has been used to evaluate noisy images caused by algae and different soil surfaces such as gritty compost, dry and wet soil types. Still, it can automatically and reliably execute the analysis tasks without users’ intervention. To verify Leaf-GP’s trait measurements, we have scored manually the key growth phenotypes on the same pots and obtained a correlation coefficient of 0.958. As the software is implemented based on open image analysis, computer vision and machine learning libraries, Leaf-GP can be easily adopted or redeveloped for other experiments. To support computational users to comprehend and share our work, we have provided very detailed comments in our source code.

From a biological perspective, the use of key growth traits generated by Leaf-GP can be an excellent tool for screening leaf growth, leaf symmetry, leaf morphogenesis and movement, e.g. phototropism. For example, the leaf skeleton is a useful tool to estimate hyponasty (curvature of the leaf). It could also be used as a marker to quantify plant maturation, e.g. *Arabidopsis* plants transits to the reproductive stage (i.e. flowering), a change from vegetative to flowering meristem when cauline leaves are produced, which can be used to mark differences in maturation. Some traits are also useful in studies other than plant development biology. For instance, vegetative greenness can be used in plant pathogen interaction to analyse the activity of pathogens on the leaf surface, as most of the times broad yellowish symptoms can be observed from susceptible plants (e.g. rust in wheat).

From a software engineering perspective, we followed best practices in computer vision and image analysis [51] when conducting feature selection, i.e. choosing traits based on the statistical variation or dispersion of a set of phenotypic data values. Whilst implementing the software, we built on our previous work in batch processing and high-throughput trait analysis [52–56] and improved software implementation in areas such as reducing computational complexity (e.g. the usage of CPU cores and memory in parallel computing), optimising data annotation and data exchange between application programming interfaces (APIs), i.e. the objects passing between internal and external functions or procedures, promoting mutual global and local feature verification (e.g. cross validating positional information of plants at the image level as well as the pot level), and implementing software modularity and reusability when packaging the software (see the software executables and package source code in https://github.com/Crop-Phenomics-Group/Leaf-GP). Furthermore, we verify that, instead of fully relying on a black-box machine learning approach without an in-depth understanding of why clustering or estimation is accomplished, it is more efficient to establish an analysis algorithm based on a sound knowledge of the biological challenge that we need to address. If the features we are interesting is countable and can be logically described, advanced computer vision and image analysis methods would be efficient for our phenotypic analysis missions.

## Conclusions

In this paper, we presented Leaf-GP, a comprehensive software application for analysing large growth image series so that multiple growth phenotypes in response to different treatments can be measured and cross-referenced over time. Our software demonstrates that treatment effects such as the response to different temperatures between genotypes could be detected reliably. We demonstrate the usefulness and high accuracy of the software based on the quantification of growth traits for *Arabidopsis* genotypes under varied temperature conditions and wheat growth in the glasshouse over time. To serve a broader plant research community, we improved the usability of the software so that it can be executed on different platforms. To help users or developers to gain an in-depth understanding of the algorithms and the software, we have provided our source code, detailed comments, software modulation strategy, and executables (.exe and .app), together with raw image data and experiment results in the Additional files. The software, source code and experiment results presented in this paper are also freely available at https://github.com/Crop-Phenomics-Group/Leaf-GP/releases.

Leaf-GP package provides an efficient and effective analysis platform for carrying out large growth phenotype measurements with no requirement on programming skills and limited requirements on imaging equipment. We followed the open software strategy so that we could share and contribute jointly with the computational biology community. Our software has confirmed previously reported results in the literature and produces a number of key growth traits that enhance the reproducibility for plant growth studies. Many plant growth and development experiments can be analysed by Leaf-GP under a range of treatment conditions. Our case studies of temperature effects and different genotypes or plant species are not limited. Natural variation in plant growth can also be analysed or images from plants experiencing mineral or nutrient stress could be equally well handled.

### List of abbreviations

RGB: a red, green and blue colour model
NoIR: no infrared filter
ROI: regions of interest
GUI: graphic user interface
HPC: high-performance computer
CSV: comma-separated values
OS: operating systems
CPU: central processing unit
Lab: lightness, a for the colour opponents green–red, and b for the colour opponents blue–yellow
RGR: relative growth rate
L*er*: Landsberg *erecta* (wildtype)
*spt-2*: spatula-2
API: application programming interfaces

## Declarations

### Ethics approval and consent to participate

Not applicable

### Consent for publication

Not applicable

## Authors' contributions

JZ, CA, NP wrote the manuscript, NP, ADA and SO performed the biological experiments under SP and SG’s supervision. JZ, NP and DR designed the plant phenotyping protocol. JZ developed and implemented the core analysis algorithm of Leaf-GP. CA, DR and MM implemented and packaged the GUI version under JZ’s supervision. JZ, CA and NP tested the software package. NP and JZ performed the data analysis. All authors read and approved the final manuscript.

## Acknowledgements

The authors would like to thank members of the Zhou laboratory for fruitful discussions. We thank Thomas Le Cornu, Danny Websdale and Jennifer McDonald for excellent technical support and research leaders at EI, JIC, TSL and UEA for constructive discussions. JZ was partially funded by BBSRC’s Designing Future Wheat Cross-institute Strategic Programme Grants (BB/P016855/1) to Professor Graham Moore. NP and ADA were supported by a Leverhulme Trust Research Project Grant (CA580-P11-H) awarded to SP. CA was partially supported by BBSRC’s FoF award (GP105-JZ1-B) to JZ.

## Authors' information

^1^Earlham Institute, Norwich Research Park, Norwich UK

^2^John Innes Centre, Norwich Research Park, Norwich UK

^3^University of East Anglia, Norwich Research Park, Norwich UK

## Availability of data and materials

All the 4.3 GB image datasets as well as The Leaf-GP software package and source code are freely available from our online repository https://github.com/Crop-Phenomics-Group/Leaf-GP/releases.

## Open Access

The software is distributed under the terms of the Creative Commons Attribution 4.0 International License (http://creativecommons.org/licenses/by/4.0/), which permits unrestricted use, distribution, and reproduction in any medium, provided you give appropriate credit to the original authors and the source, provide a link to the Creative Commons license, and indicate if changes were made. Unless otherwise stated The Creative Commons Public Domain Dedication waiver applies to the data and results made available in this paper.

## Additional files

- **Additional File 1**: The interactive *Jupyter Notebook version* for Leaf-GP (version 1.18)
- **Additional File 2**: Install manual for Python environment, Anaconda Python distribution and OpenCV-Python binding
- **Additional File 3**: Processed images of *Arabidopsis* rosettes at different growth stages
- **Additional File 4**: Multiple trait measurements results based on a testing series
- **Additional File 5**: The analysis workflow and a detailed activity diagram of Leaf-GP
- **Additional File 6**: The manual for importing image datasets via the GUI version of Leaf-GP
- **Additional File 7**: The *Jupyter Notebook version* for plotting and cross-referencing growth traits between experiments
- **Additional File 8**: Multiple trait measurements results based on L*er* 12°C, 17°C, 22°C
- **Additional File 9**: Multiple trait measurements results based on *spt-2* 12°C, 17°C, 22°C

**Figure S1.**
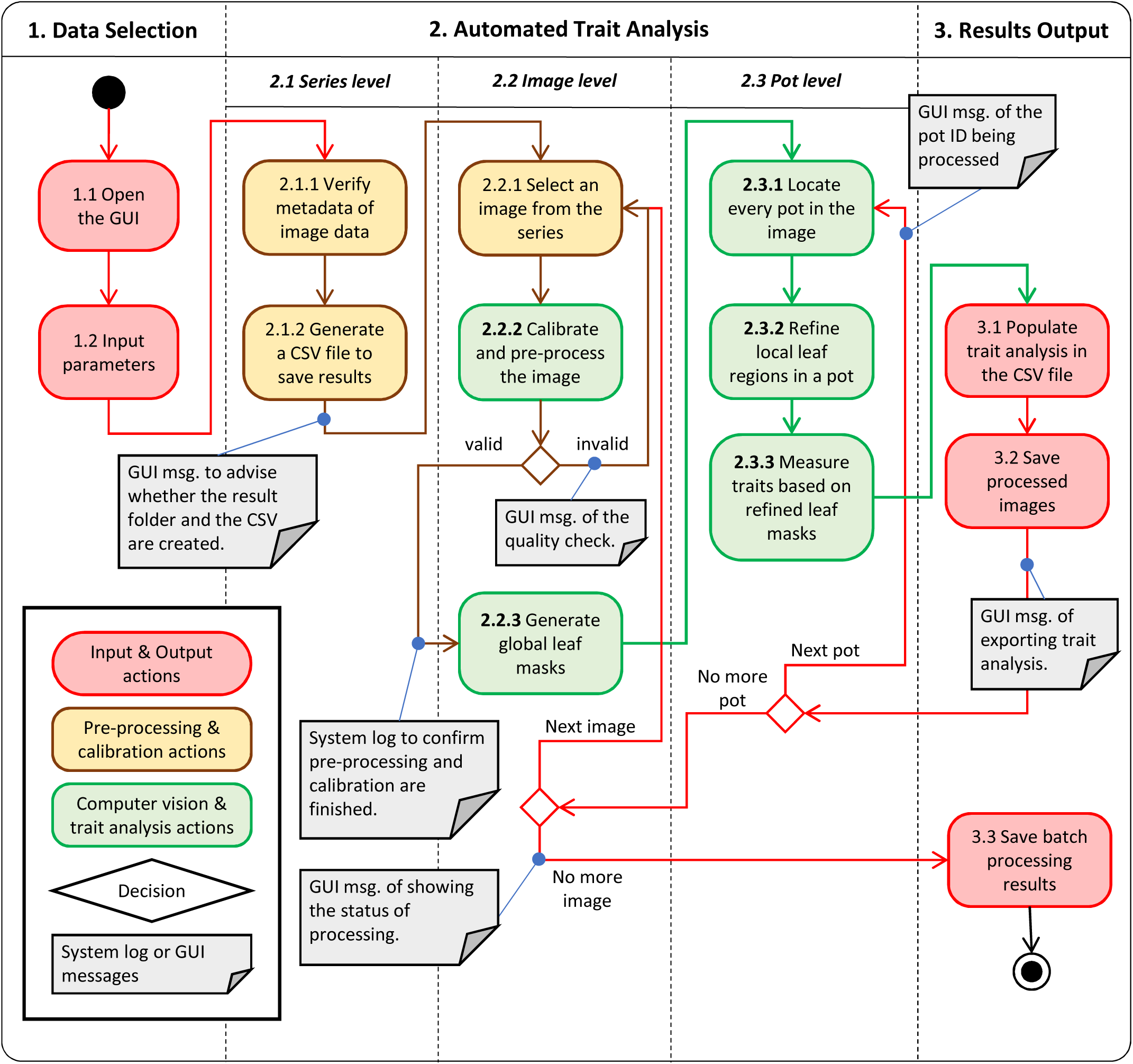
The Analysis Workflow of Leaf-GP

**Figure S2.**
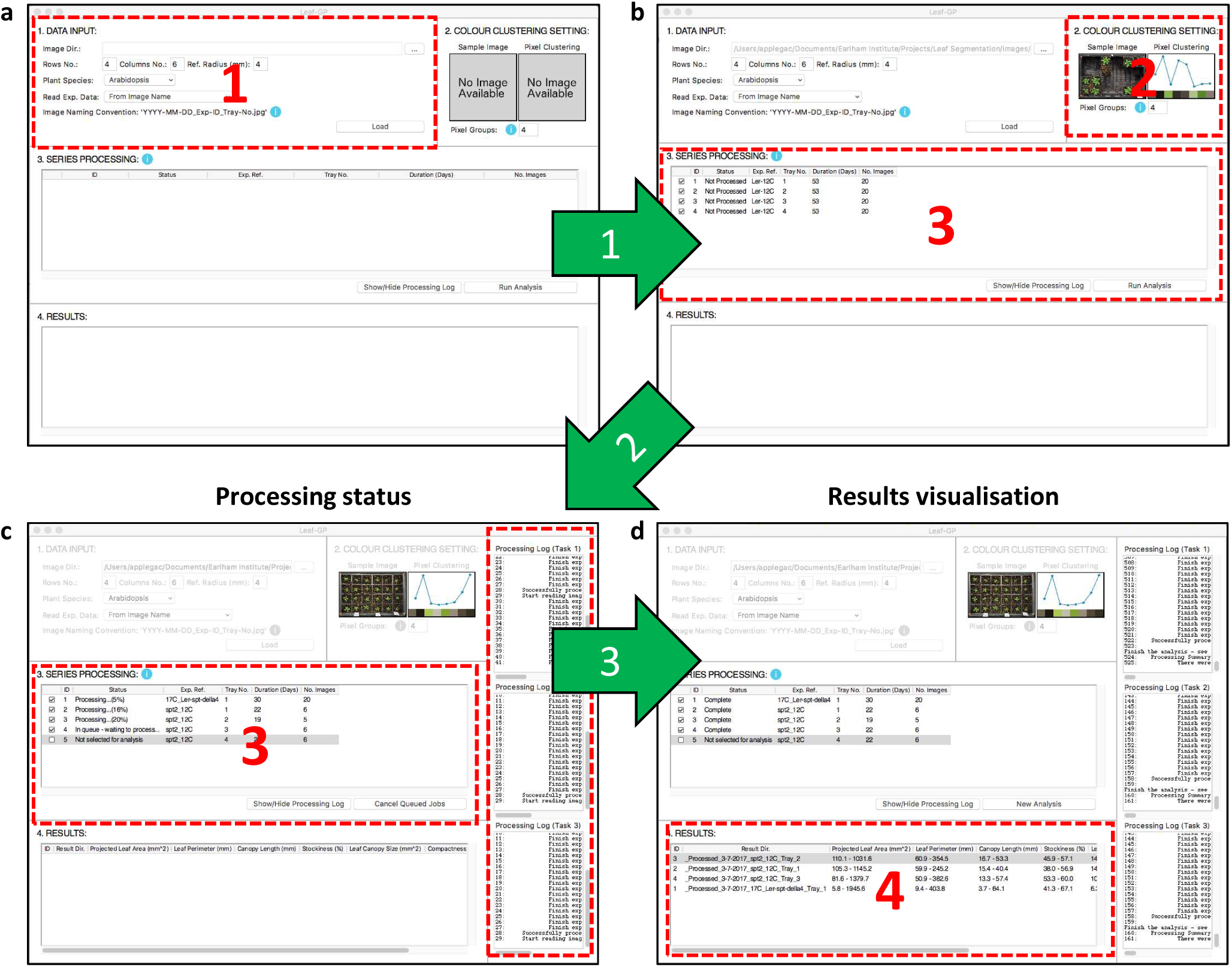
The GUI operation

